# A chromosome-level genome assembly and resequencing data reveal low DNA methylation and reduced diversity in the solitary bee pollinator *Osmia cornuta*

**DOI:** 10.1101/2025.07.14.664667

**Authors:** Jannik S. Möllmann, Xuejing Hu, Eva A. Baumgarten, Anne Hartleib, Katja Nowick, Thomas J. Colgan, Eckart Stolle

## Abstract

Bees provide essential pollination services that contribute to ecosystem stability, as well as the sustainability of economic crop yields. Due to concerns over global and local declines, improving our understanding of these ecologically and commercially important species is crucial for determining their capacity to respond and adapt to environmental challenges. The European orchard bee (*Osmia cornuta*) is a solitary bee of increasing agricultural importance due to its role in the pollination of fruit crops, yet lacks genomic resources. Using cost-effective Nanopore-only long-read sequencing, we report the first genome assembly for *O. cornuta*, spanning 647.56Mb across 727 contigs (N50=3.94Mb) at a high level of completeness (99.88% BUSCO complete). In line with the expected number of chromosomes in this species, 16 major scaffolds were assembled to chromosome-level. Also, we provisionally investigated the epigenomic architecture of *O. cornuta*, finding low numbers of CG dinucleotides that were either 5’-methylated or 5’-hydroxymethylated, providing additional evidence for the limited role methylation plays in gene regulation in Hymenopterans. To generate improved gene annotations, we combined transcriptomic- and orthology-based approaches, leading to the prediction of 12,144 genes and 25,964 proteins, showing exceptionally high BUSCO completeness (99.64%). Lastly, through whole-genome resequencing of a representative dataset, we provisionally find patterns of reduced nucleotide diversity and lower recombination rates within *O. cornuta* compared to other bee species. Collectively, our study provides a novel insight into the genome architecture of a key pollinator, providing an important resource to facilitate further genomic studies.

**Significance statement:** Solitary bees are crucial pollinators for crops and wild plants, but scientists have lacked the genetic blueprints needed to understand how these species might adapt to environmental threats like climate change and habitat loss. Using modern and cost-effective technologies, we generated the complete genome of the European orchard bee, an economically important fruit pollinator. This comprehensive genetic resource provides the foundation for future research on solitary bee biology and conservation, necessary to understand their adaptive potential in the face of environmental challenges. In line with this, we analysed this species’ genetic variation across populations in Germany using a small, representative dataset and found surprisingly low genetic diversity and limited evidence of gene recombination during reproduction, which are factors that could restrict its capacity to evolve in response to environmental challenges.

## Introduction

Improving our fundamental understanding of the biology of ecologically and commercially important pollinators is particularly important considering they provide key ecosystem services that are crucial for ecosystem stability, biodiversity, and sustainability of agricultural yields (Kremen et al. 2007; Potts et al. 2016; Gallai et al. 2009). Despite this, declines in natural populations have raised concerns about the long-term viability of such pollination services (Hallmann et al. 2017; Powney et al. 2019; Lippert, Feuerbacher, and Narjes 2021). Key factors that are proposed to contribute to declines include habitat loss and fragmentation, pesticides, parasites and pathogens, invasive species, and changes in climatic conditions (Cameron et al. 2011; Potts et al. 2016; Seibold et al. 2019; Möllmann et al. 2024). In addition, for certain pollinators, such as solitary bees, the lack of social buffering commonly found in social bee colonies may mean they are predicted to be even more susceptible to environmental challenges that contribute to declines (Straub et al. 2015). It is, thus, of vital importance to improve our understanding of the biology of solitary bees to inform on their capacity to respond and adapt to environmental challenges.

An important solitary bee of ecological and commercial importance is the European orchard bee *Osmia cornuta* (Latreille, 1805), which is native to most of Central and Southern Europe with its range extending south to Northern Africa and east to Central Asia (Banaszak and Romasenko 1998). It belongs to a taxon-rich genus of over 300 species (Praz et al. 2008), which are part of the family Megachilidae. While the commercial use and supply of mason bees, such as *O. cornuta*, due to their pollination efficiency, especially in apple orchards, has led to their increased presence in scientific studies, they also contain certain traits, such as its univoltine life-cycle, generalist foraging patterns, and nesting behaviour (Splitt, Schulz, and Skórka 2022), which make *O. cornuta,* an increasingly suitable model to study and understand aspects that are more common to the biology of solitary bees (Jordi Bosch and Vicens 2006; Vicens and Bosch 2000; J. Bosch 1994; Jordi Bosch and Kemp 2004; Fauria and Campan 1998; Strobl et al. 2019). *O. cornuta* has also raised recent ecological concerns over its impact through accidental introductions into locations where it is non-native. For example, due to its efficiency as a pollinator of fruit crops (Gruber et al. 2011; Sedivy and Dorn 2014; Fliszkiewicz and Giejdasz 2023), cocoons of *O. cornuta* have been traded to North America and other places outside of Europe where recent reports have confirmed the presence of feral populations that have begun to establish in British Columbia, Canada (Getz et al. 2024), likely as a consequence of trade.

Recent advances in sequencing technologies and the accompanying development of genomic tools to analyse these data have facilitated the rapid generation of genetic data that has informed on the genomic architecture of non-model organisms, including solitary bees (Möllmann and Colgan 2022; Beadle et al. 2019; Barribeau et al. 2015; Kapheim et al. 2015, 2019; Santos et al. 2025; R. Wang et al. 2025; Stolle et al. 2023; Yang et al. 2025), which has the potential for application in the fields of agriculture, as well as conservation and wildlife management approaches (Allendorf, Hohenlohe, and Luikart 2010; Kelemen and Rehan 2021; Trapp, McAfee, and Foster 2017). For example, through the generation of whole-genome sequencing data, it is now possible to accurately determine millions of nucleotide bases providing a solid basis to allow for estimating genetic parameters relevant for ecological modelling and conservation biology, such as genetic variation, effective population size and population connectivity, or identify regions of the genome with signatures of selection, helping to understand the adaptive potential of these species (Webster et al. 2023; Eizaguirre and Baltazar-Soares 2014). Similarly, transcriptomic-based methods enable the inference of accurate gene models as the basis for the identification of candidate genes and molecular pathways involved in responses to environmental stressors, such as climate change and pesticide exposure (Beadle et al. 2019; Möllmann and Colgan 2022).

In order to leverage these approaches, however, the availability of a high-quality chromosome-level genome assembly, including accurately predicted gene models, is of utmost importance as it underpins all genetic and transcriptomic analyses performed. For a few members of the genus *Osmia*, genome assemblies are already available (Beadle et al. 2019), however, some of these were assembled using short-read data, meaning they are likely less contiguous and more fragmentary, and may suffer from lower quality gene model predictions, limiting their utility compared to long-read generated chromosome assemblies (R. Wang et al. 2025; Ouyang et al. 2025). Therefore, for *Osmia* species, such as *O. cornuta*, improvements in the quality of genome assembly, which can be generated to chromosome-level using long-read sequencing data, are urgently warranted to improve our understanding of the genetic architecture of these species, as well as provide sufficient resources to profile the genetic variation harboured in natural populations and resolve aspects of their recent demographic history. Here, we present a genome-based analysis of *O. cornuta*, for which we sequenced an individual male to high sequencing depth using Oxford Nanopore long-read sequencing technology, which we subsequently assembled to chromosome-level and generated high-quality gene model predictions using a combination of transcriptomic- and orthology-based gene annotation approaches. We also provide insights into the epigenomic landscape of *O. cornuta* through examination of DNA methylation levels, genome complexity through analysis of repeat diversity, as well as perform provisional analyses of nucleotide diversity and recombination rates through whole-genome sequencing of individuals sampled from a European population.

## Results and Discussion

### High-completeness, chromosome-level genome assembly from long-read sequencing data

From a single *Osmia cornuta* male (Figure 1A), we generated 1,338,458 highly accurate, filtered reads on approximately half of a Nanopore PromethION flow cell at a total length of 26.12Gb (mean length = 19.52kb; maximum length = 930.51kb). Filtered reads were then corrected using HERRO, resulting in 1,110,145 reads at a total length of 20.94Gb (mean length = 18.86kb; maximum read length = 139.28kb). Using these reads, we generated 11 candidate assemblies, six of which were generated using flye and five were generated using hifiasm. Based on variation in contiguity, completeness, and the presence and extent of telomeres, we selected an assembly generated using hifiasm for further research and publication. This assembly initially contained 777 contigs (total length = 654.189Mb), but was reduced by 44 contigs following purging of putative duplicate haplotigs. It had an excellent merqury QV score of 54.86, corresponding to a very low error rate (estimated error rate = 1 error per 306,000 bases), indicative of a high-quality, highly accurate assembly. Similarly, screens of foreign DNA did not classify any contigs as being contaminants. A scan for mitochondrial sequences suggested that six contigs were putative mitochondrial in origin. After removal of mitochondrial-derived contigs, the final assembly contained 727 contigs, without gaps, and a total length of 647,555,333bp (Figure 1B) with a BUSCO completeness score of 99.78%. The longest contig spanned 21.64Mb, the mean length of contigs was 890.72kb and N50, N70, and N90 lengths were 3.94Mb (n=29 contigs), 1.35Mb (n=91) and 0.34Mb (n=271), respectively. By mapping the reads back to the genome assembly, we inferred a mean coverage of 40.22X (SD=32.14), whereas less repetitive parts of the largest primary contigs (n=16) showed a coverage of approximately 55.14X. The high completeness of our assembly was further supported by the identification of 15 complete telomeres (AACCT) among the 16 largest contigs, which represent the 16 putative chromosomes previously described via karyotyping (Kerr and Laidlaw 1956). Another nine putative telomeres were found on shorter (<1.5Mb) contigs. Additionally, we assembled the mitochondrion contig to a length of 35,525bp, containing the expected two ribosomal RNA (rRNA), 24 transfer RNA (tRNA), and 13 protein-coding genes. A comprehensive annotation of repetitive elements determined that 80.3% of the genome was composed of repeats (Figure 2C), of which the majority were LTR retrotransposons (34.29% of the assembly) and simple repeats (29.78%), while other types were much less common (DNA transposons: 7.25%, LINE: 6.67%, Rolling Circle: 0.6%, and Unclassified Repeats 1.7%). Together, the identification of the presence and extent of telomeric regions and the exceedingly high BUSCO completeness suggest a highly contiguous genome assembly, comparable with high-quality PacBio and Hi-C genome assemblies of other bee species. (Toth et al. 2024; Wallberg et al. 2019; R. Wang et al. 2025; Stolle et al. 2023)

**Figure 1.**
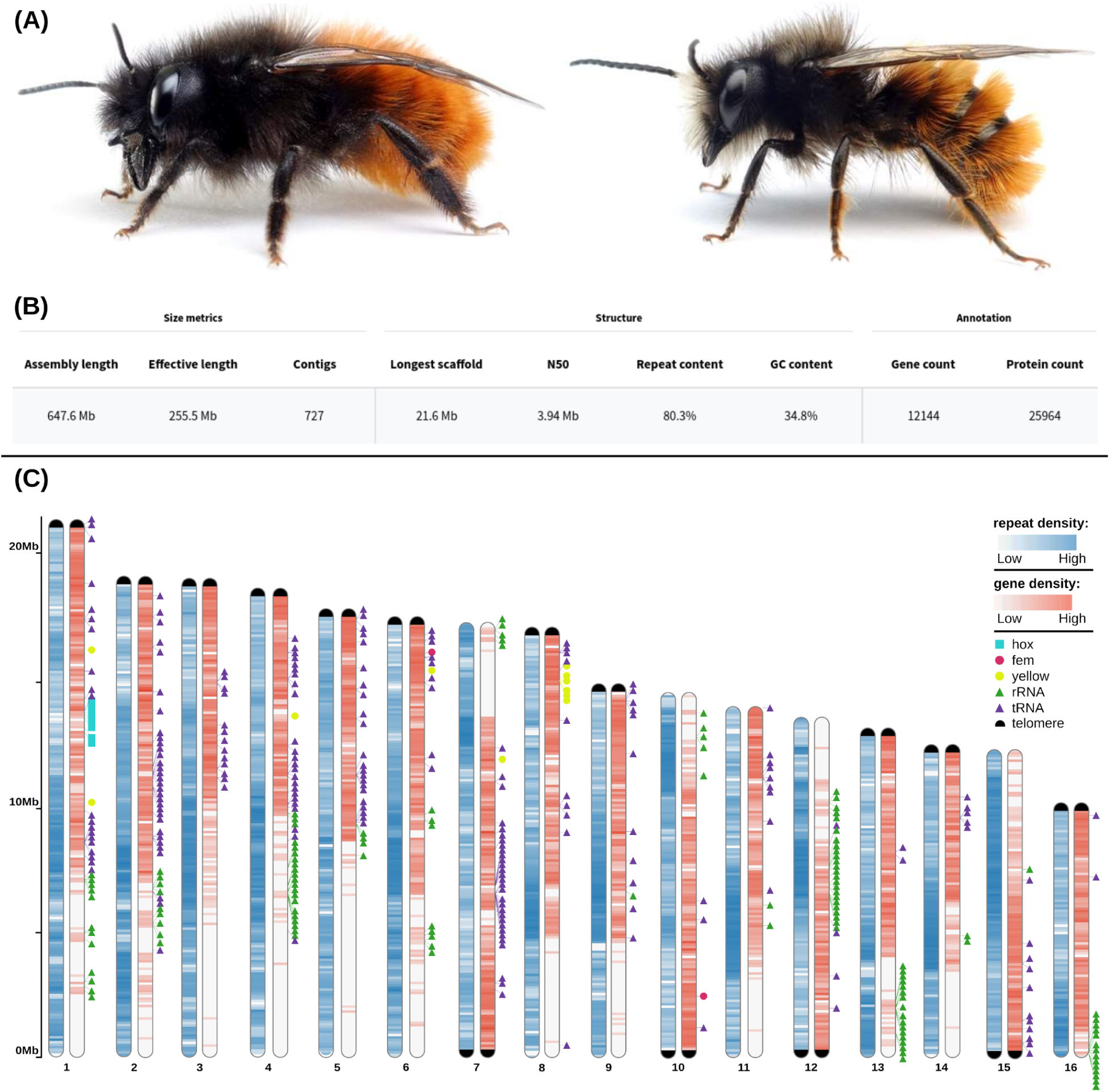
Habitus, overview statistics and chromosomal karyotype. **(A)** Habitus of *Osmia cornuta*, depicting both a female (left) as well as a male (right) individual (image copyright: www.apidarium.de). **(B)** Assembly and annotation summary statistics, classified into size-related, assembly structure-related, and annotation-related metrics. **(C)** A chromosomal karyotype that depicts for each of the 16 chromosomes regions of increased or reduced gene- and repeat-density. In addition, the presence and location of some representative gene families, which tend to cluster in specific regions of the genome, are depicted using coloured symbols. An additional, black symbol is used to mark telomeres at chromosomal ends. The size of this symbol is uniform and does not represent the actual length of these telomeric regions.

**Figure 2.**
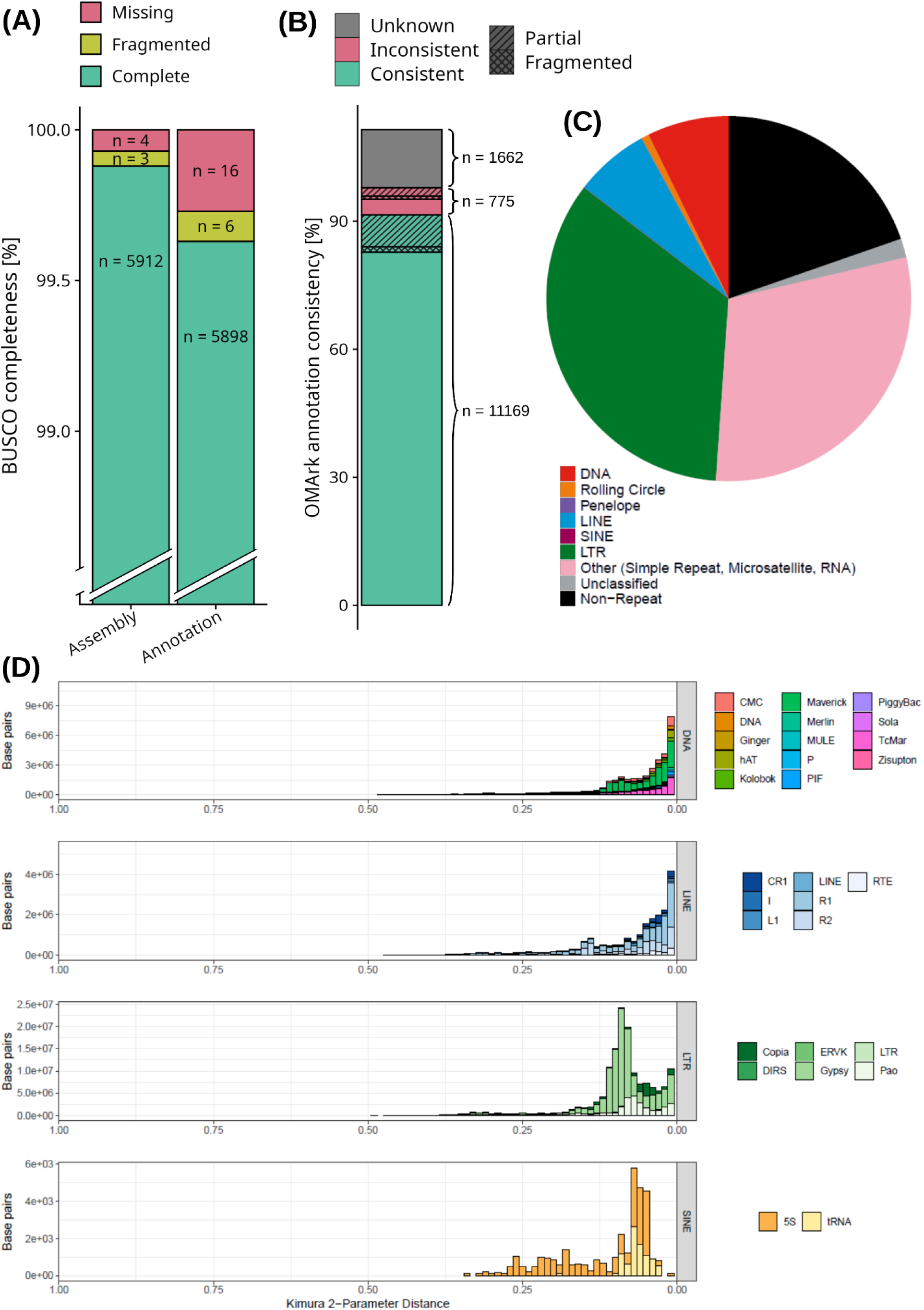
High assembly- and annotation completeness and overview of genomic repeat architecture. **(A)** Stacked bar charts showing the results of the overlap of the assembly (left) and the predicted protein models of the annotation (right) with single-copy orthologs from the hymenopteran dataset (n=5919) of the OrthoDB (v12) database. Within each bar chart, the level of overlap for three subcategories is colour-coded: “Complete”, green, refers to an assembly region or a protein fully overlapping with an ortholog in the reference database; “Fragmented”, yellow, refers to an assembly region or a protein that partially overlaps with a reference ortholog; and “Missing”, red, which refers to a reference ortholog that did not overlap with any region of the assembly or predicted protein set. **(B)** A bar chart showing the annotation consistency of the predicted gene models when compared against hierarchical orthogroups (HOGs) of the OMA database with subcategories either colour-coded or patterned: “Consistent”, green, referring to a gene model has been placed in the correct lineage; “Inconsistent”, red, referring to a gene model that has been placed in an incorrect, unspecific lineage; “Unknown”, gray, which refers to a gene model has not been placed in any of the tested lineages; “Partial”, stripes, meaning not all parts of a gene model match a HOG; and “Fragmented”, checkered, meaning a gene model does not match all parts of a HOG. **(C)** A pie chart showing the genomic composition by repeat type and the fraction of non-repeat regions. Collectively, more than three quarters of the genome are made up of repeat sequences with long terminal repeats (LTR) being the most common. Legend: DNA = DNA transposon; Rolling Circle = rolling circle transposon; Penelope = Penelope transposon; SINE = short interspersed nuclear element; LTR = long terminal repeat; and RNA = retrotransposon. **(D)** Four histograms, each depicting the distribution of the kimura 2-parameter distance in base pairs for a specific type of repeat. Within each histogram, the repeat type is broken up into subtypes, each displayed at a different color.

### High-quality structural annotations generated using a combination of protein-, transcript- and orthology-based evidence

We generated gene model predictions for the assembly using two independent approaches: first, using transcript- and protein-based evidence (implemented with BRAKER3); and second, using inferred orthology with the closely related *O. bicornis* (implemented with TOGA). We then combined predictions from these two approaches to generate a more complete set of gene models. The combined set of predictions comprised 12,144 genes predicted to encode for 25,946 protein-coding transcripts. The resulting protein-coding transcript predictions showed a BUSCO completeness score of 99.64%, whereas proteins from the individual annotation approaches resulted in completeness scores of 97.20% and 97.51%, respectively. Compared to the completeness score of 99.88% for the assembly (Figure 2A), our high completeness score for the annotation suggests that, using this combined approach, we were able to predict gene models for nearly all genes that were assembled. The high completeness of the structural annotations was further highlighted by a high percentage (97.64%) of matches against conserved and taxonomically ‘consistent’ hierarchical orthogroups (HOGs) from the Orthologous Matrix (OMA) database, as well as a low percentage (3.63%) of matches against HOGs that are considered taxonomically ‘inconsistent’ with the species of interest (Figure 2B). These numbers are highly comparable to the structural annotation of the closely related *O. bicornis* (<25 million years of divergence (Cardinal and Danforth 2013; Gonzalez 2011), 97.68% completeness, 3.67% ‘inconsistency’ of proteome), which has been annotated via the NCBI Eukaryotic Genome annotation pipeline, indicating that our structural annotation has achieved a similar level of quality. In line with the high completeness of our gene annotations, we successfully confirmed the presence of several marked gene clusters and genes (Figure 1C), among which the *yellow* (chromosome 8), *hox* (chromosome 1) and *osiris* clusters (chromosome 4), as well as the *feminizer* gene (chromosome 6), which forms part of the sex determining cascade. As our annotation approach did not predict non-coding gene models, we additionally scanned the chromosomal scaffolds of our assembly for tRNA and rRNA genes. In doing so, we found 195 tRNA genes with particularly pronounced clusters on chromosomes 2, 4, and 7, as well as 126 rRNA genes with strong clustering on chromosomes 4, 12, 13, and 16. Lastly, we also examined synteny of our genome assembly with that of the closely related *O. bicornis* via comparison of predicted proteomes. Summarised to gene level, we found that 77.43% of genes were collinear between the two species. Using this synteny-based approach, we further identified 16 chromosomal homologues to those described in *O. bicornis*, providing evidence of the high quality and completeness of the *O. cornuta* genome assembly. Additionally, we also found evidence of inverted segments on several chromosomes within the *O. cornuta* genome assembly, reflecting putative chromosomal rearrangements since the latest common ancestor (Figure 3). In summary, our combined annotation approach produced a highly complete set of gene predictions for *O. cornuta*, successfully identifying key gene clusters, and demonstrating strong syntenic conservation with the closely related *O. bicornis*.

**Figure 3.**
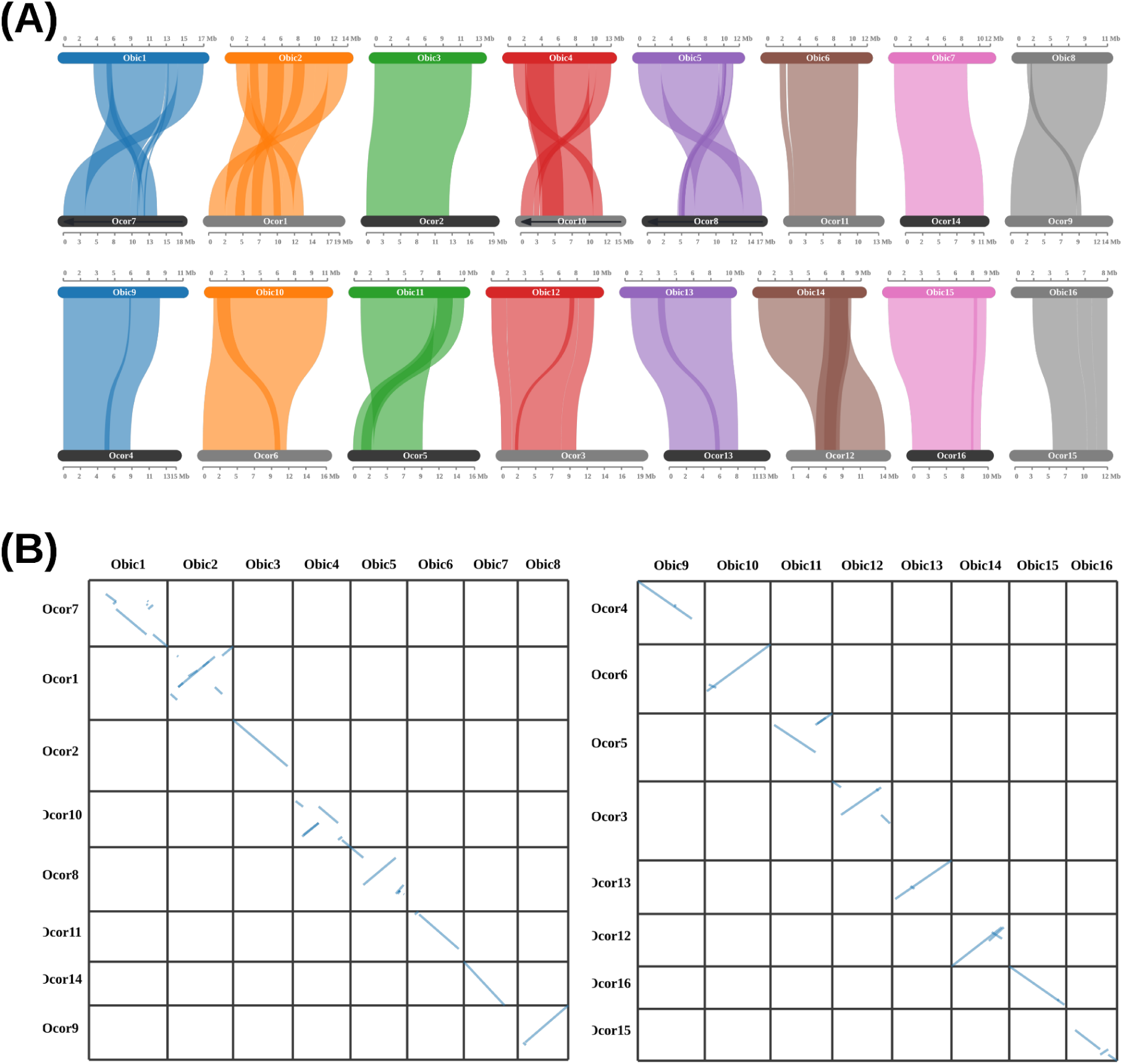
Synteny comparison with *Osmia bicornis*. **(A)** A two-row parallel link plot giving an overview of syntenic regions between putative chromosomal scaffolds of *O. bicornis* (Obic; top of each row) and *O. cornuta* (Ocor: bottom of each row) based upon comparisons of predicted proteomes. The markers show the genomic coordinates for each putative chromosome in megabases (Mb). **(B)** Dot plots displaying a more detailed examination of synteny between the two species. The binned x-axis represents the position along identified chromosomes in *O. bicornis* with one chromosome per bin. Similarly, the y-axis represents the position along chromosomes in *O. cornuta* with one chromosome per bin. A dot represents a syntenic region that is shared between the two species at the respective chromosomal loci.

### Evidence of genome-wide methylation patterns

A key advantage of Nanopore technology is its ability to detect modified bases, such as methylated cytosine, during sequencing, allowing for a direct examination of the epigenomic landscape. Methylation levels have previously been described as low in members of the Hymenoptera, including other bee species, with studies employing bisulphite sequencing, identifying approximately 1% of genome-wide cytosines reported as methylated in the honeybee *A. mellifera* ((Harris et al. 2019), and the bumblebee *B. terrestris* (Pozo et al. 2021). We, therefore, examined whether the genome of *O. cornuta* contained similar levels of methylated cytosines. By extracting base modification information directly from Nanopore data through basecalling, we found that the *O. cornuta* genome assembly contained no 6’-methyladenosine (6mA), 5’-methylcytosine (5mC), or 5’-hydroxymethylcytosine (5hmC) modifications, but low numbers of 5’-methylcytosine and 5’-hydroxymethylcytosine in the context of CG dinucleotides (5mCG, 5hmCG). For 5mCG, out of 24,194,970 called sites, 192,079 (0.84%) were considered methylated, being supported by at least one read, while 51,437 (0.23%) sites showed methylation within more than 20% of reads. For 5hmCG, out of 24,194,970 called sites, we identified 201,054 (0.9%) sites that were considered methylated, being supported by at least one read, while 14,125 sites (0.06%) showed methylation in more than 20% of reads. To understand how 5’-methylation of CpG loci varied across different regions and features of the genome, we inferred for each investigated genomic feature type, the percentage of sites that were considered weakly methylated, finding evidence of considerable variation in methylation patterns across the genome (Table 1). In particular, exons showed by far the highest proportion of 5’-methylated CpG sites (2.53%), while introns and intergenic regions had a relatively lower proportion of 5’-methylated CpG sites (intron: 0.88%, intergenic: 1.02%). Strikingly, in low complexity and repeat regions, the frequency of 5’-methylated CpG sites was up to 100x reduced compared to all other genomic feature types (low complexity: 0.01%; repeats: 0.03%). The same overall patterns were observed for 5’-hydroxymethylcytosine CpG modifications. Collectively, we found that DNA methylation levels within the genome of *O. cornuta* are low, but comparable to levels previously reported for other hymenopterans, providing additional support for the growing literature highlighting the likely reduced role of DNA methylation within gene regulation for these species.

**Table 1.** Genome-wide patterns of methylation across different genomic features. A table showing for each genomic feature considered in the analysis, the percentage as well as the number of CpG sites for which we found evidence of either a 5’-methylcytosine modification (5mCG) or 5’-hydroxymethylcytosine (5hmCG) modification. In addition, the total number of potentially methylated CpG sites for each genomic feature is shown in a separate column.

| Genomic Feature | 5mCG |  |  | 5hmCG |  |  |
| --- | --- | --- | --- | --- | --- | --- |
|  | Methylated sites (%) | Methylated Sites (n) | Methylatable Sites (n) | Methylated Sites (%) | Methylated Sites (n) | Methylatable Sites (n) |
| Exon | 2.53 | 23,738 | 936,051 | 1.79 | 16,781 | 936,051 |
| Intron | 0.88 | 37,240 | 4,190,504 | 1.11 | 46,615 | 4,190,504 |
| Intergenic | 1.02 | 83,839 | 8,198,788 | 1.17 | 95,525 | 8,198,788 |
| Low Complexity | 0.01 | 8,654 | 7,448,992 | 0.02 | 12,157 | 7,448,992 |
| Repeat | 0.03 | 54,052 | 15,547,674 | 0.06 | 89,611 | 15,547,674 |

### Assessment of intraspecific genetic variation informed by whole-genome resequencing

As a preliminary examination of the genetic variation maintained in *O. cornuta*, we generated whole-genome resequencing data for 10 females sampled from across Germany (sampling sites in Supplementary Table 1). We identified 594,857 high-quality, filtered SNPs (biallelic: 99.7%) across the 16 chromosomal scaffolds of our assembly (255.5Mb effective length), yielding an overall variant density of 2.33 SNPs per kilobase. Variant density varied across chromosomal scaffolds, ranging from 1.73 (Chromosome 3) to 3.22 SNPs per kilobase (Chromosome 15). Structural annotation revealed that the majority of variants (94.9%) were positioned in non-coding regions, such as intronic, intergenic, and regions upstream/downstream of genes. Among protein-coding regions, 74.5% of variants (n=22,116) were synonymous (silent) polymorphisms preserving amino acid sequences, while 25.3% (n=7,517) were nonsynonymous (missense) variants, predicted to alter the amino acid sequence of the resulting protein. Protein-truncating variants, such as early stop codons, were rare, with nonsense mutations attributed to only 0.14% (n=42) of variants within protein-coding regions and critical splice site disruptions caused by only 21 variants genome-wide. The overall missense-to-silent ratio of 0.34 likely reflects the expected purifying selection acting against amino acid-altering mutations.

### Reduced genetic diversity and low recombination rates

Using our set of called variants, we calculated genome-wide estimates of nucleotide diversity in non-overlapping 10kb-sized windows finding a median diversity of *π*=0.0012 (Figure 4A) as well as a median Watterson’s theta of *θ_w_*=0.001 across all 16 chromosomal scaffolds. This is considerably lower than whole-genome sequencing based published estimates of nucleotide diversity in eusocial species of the genera *Apis* and *Bombus (Leroy et al. 2024; Heraghty, Jackson, and Lozier 2023; Thomas J. Colgan et al. 2022; Yinchen Wang et al. 2024)* which range from *π*=0.0014 to 0.0025 but is also lower than the estimate inferred for another solitary species of the same family (Megachilidae: *Megachile rotundata*) at *θ_w_*=0.003 (Jones et al. 2019). This pattern is likely reflective of our lower sample size, which likely captures less genetic variation compared to other studies. We also identified heterogeneity in the chromosomal distributions of nucleotide diversity (Figure 4B). Two chromosomal scaffolds (9 and 15) showed significant overrepresentation among the top 5% of windows in terms of elevated nucleotide diversity compared to the genome-wide background (Fisher’s exact test, Benjamini-Hochberg adjusted *p* < 0.01). We also assessed nucleotide diversity differences across specific genomic regions and found that median nucleotide diversity was significantly elevated in both intronic (*π=*0.0017) and repeat regions (*π=*0.0016) compared to the rest of the genome (*π=*0.0012, Wilcoxon rank-sum test, *p* < 1×10⁻¹⁶), but was not significantly elevated or decreased in intergenic (*π=*0.0012) or low complexity (*π=*0.0014) regions. In contrast, exonic regions (*π=*0.0006), were significantly decreased in diversity compared to the rest of the genome (Wilcoxon rank-sum test, *p* < 1×10⁻¹⁶), which, especially compared to the three times higher diversity estimates found in introns, may be interpreted as potential evidence of different selective pressures acting on coding and non-coding regions of the genome (Figure 4A). Linkage disequilibrium (LD) analysis revealed comparatively slow LD decay across the genome (Figure 4C). The correlation coefficient (r²) stayed at values above 0.3 for distances below 100bp and converged to around 0.1 by 10kb of distance separating linked loci, whereas in another study on another species from the same family (Megachilidae: *M. rotundata*) LD decay reached values as low as around 0.05 by 10kb of distance (Jones et al. 2019). In the same study, the recombination rate for *M. rotundata* was reported to be 1.02cM/Mb. In contrast, the advanced eusocial western honey bee (Apidae: *Apis mellifera*) reached LD levels as low as 0.05 already at distances below 100bp due to its extremely high recombination rate of 23.94cM/Mb (Jones et al. 2019). Together, our preliminary assessment provides evidence of reduced nucleotide diversity, albeit with peaks of elevated genetic diversity (Fig. 4), and relatively low rates of recombination both at shorter and longer physical distances in *O. cornuta* compared to social honeybees (23.94cM/Mb), bumblebees (4.76-8.7cM/Mb), stingless bees (9.3–12.5cM/Mb) and solitary leafcutter bees (1.02cM/Mb) (Liu et al. 2017; Waiker et al. 2021; Stolle et al. 2011; Jones et al. 2019).

**Figure 4.**
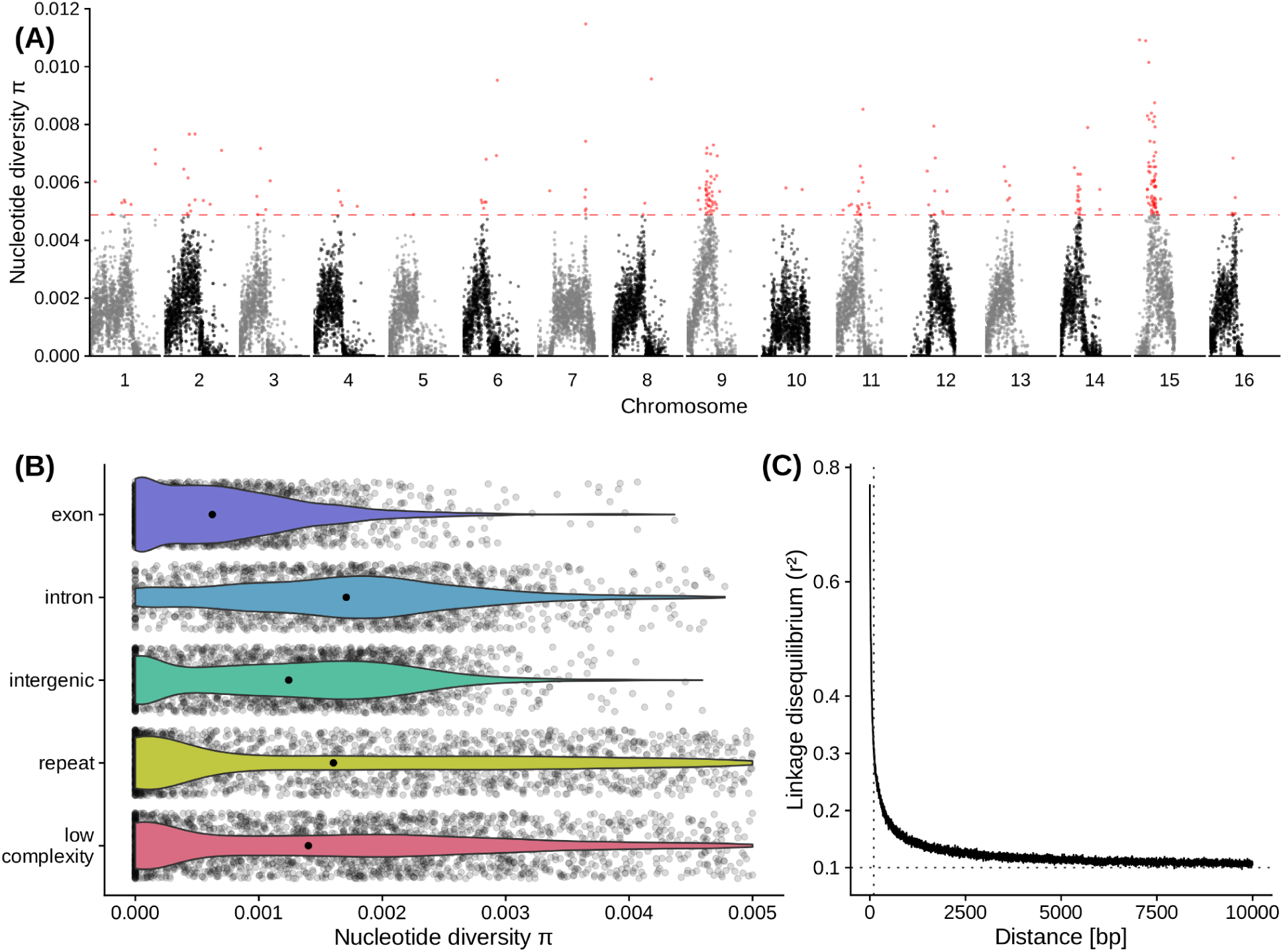
Low nucleotide diversity and low linkage disequilibrium decay inferred from whole-genome-resequencing. **(A)** A Manhattan plot showing the distribution of nucleotide diversity estimates, as measured in π, along each of the 16 chromosomes of the *O. cornuta* genome assembly. To achieve a higher resolution, each dot represents a 10kb window instead of the 100kb window that we used to estimate the values reported in the text. Coloured dots above the horizontal dashed line represent the top 5% most diverse windows across the whole-genome. **(B)** Violin plots showing the distributions of nucleotide diversity estimates across primary genomic feature types with each feature type on a separate row and indicated by an individual color. The single black dots in the middle of each violin plot mark the median while transparent dots in the background show the raw data points (10kb windows). **(C)** A line graph showing the logarithmic decay of linkage disequilibrium (y-axis) with increasing distance between loci (x-axis). Linkage disequilibrium was computed in terms of r². A dotted horizontal line marks the value of 0.1, which is the value of convergence for this curve, whereas a vertical dotted line marks a distance of 100bp that can be used to compare LD decay across species. Here, at 100bp, LD has decayed to a value of around 0.3.

### Reduced effective population size around the last glacial maximum

From our WGS data, we inferred historic effective population sizes for *O. cornuta*, which suggest a decline towards and after the last glacial maximum (Fig. 5). For more recent historic times, the effective population sizes were consistent for samples between 10,000 and 30,000 with exception of two samples. This species showed a rapid northward expansion in the last 10 years in central Europe, not yet reflected in this inference. Since rapidly expanding populations are often characterized by lower genetic diversity near their expansion wave front, our current sampling may underestimate the genetic diversity of *O. cornuta* and, hence, bias the inference of historic effective population sizes. Consequently, a re-evaluation is recommended with more data and samples to confirm the observed patterns.

**Figure 5.**
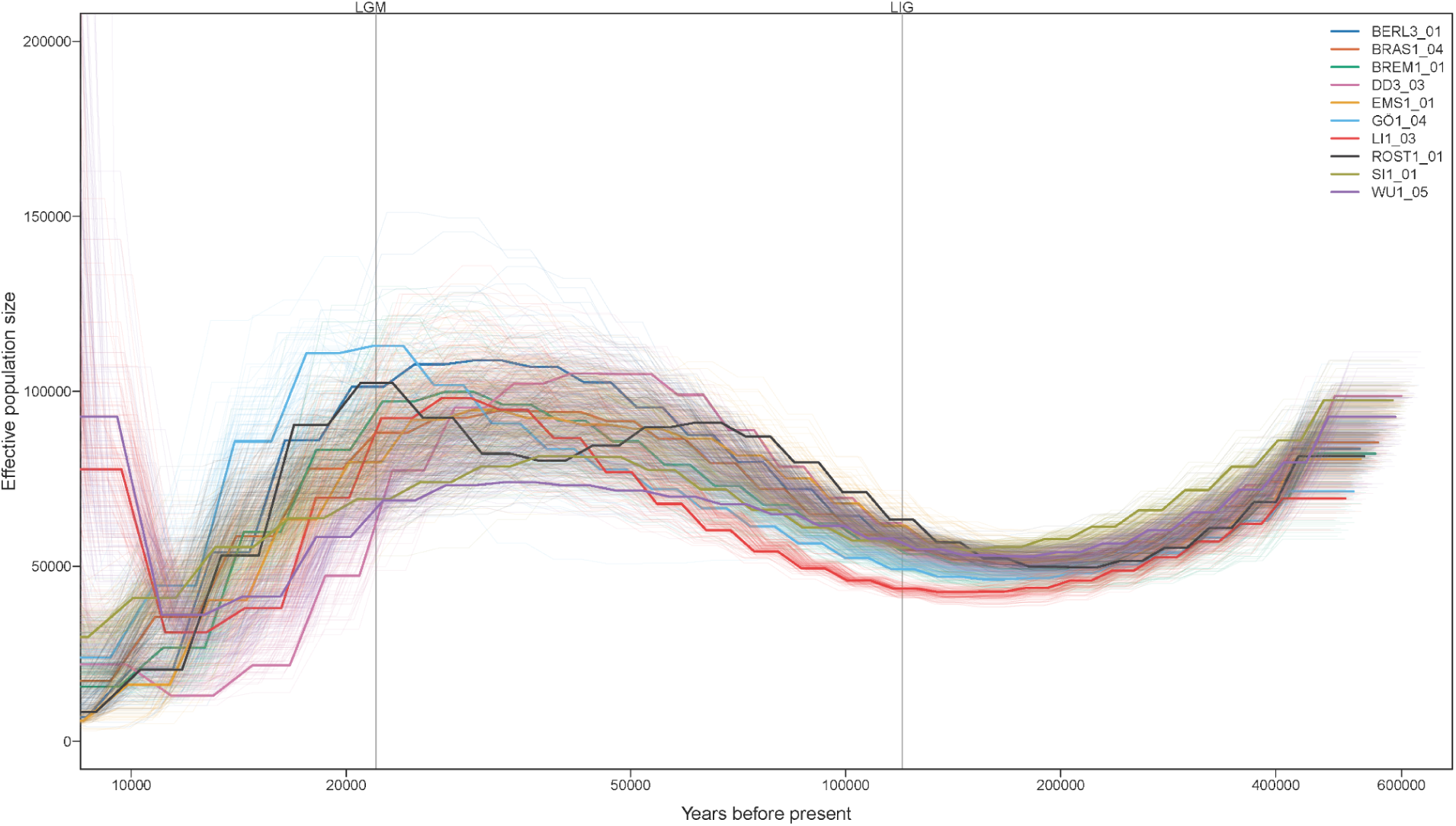
Effective population size is estimated to have declined since the Last Glacial Maximum. Line plot showing inferred demographic history of ten female *O. cornuta* sampled across Germany. The y-axis represents effective population size, while the x-axis indicates time in years before present. Each individual is represented by a distinct colour, with faint lines illustrating the variation in effective population size estimates based on 100 bootstrap iterations. Vertical grey lines correspond to the Last Glacial Maximum (LGM) and Last Interglacial (LIG) period, respectively.

## Conclusion

Our study presents a highly contiguous, chromosome-level genome assembly, complete with a high-quality genome annotation for the ecologically and commercially important solitary bee *O. cornuta*. Given the lack of genomic resources that exist for *O. cornuta*, as well as other mason bees, this genome assembly provides a valuable resource for future genetic and genomic analyses to inform on how evolutionary processes are shaping their capacity to adapt to environmental pressures. We provide a demonstration of this potential by detailing preliminary measures of genetic variation, recombination rates, genome-wide methylation, as well as demographic changes within a European population. Our project also demonstrates the feasibility to generate chromosome-level assemblies of high completeness *and* genome-wide DNA methylation profiles using as little as half of an Oxford Nanopore PromethION flow cell, which has an estimated cost of ∼€500, providing growing support to how improvements in sequencing costs and throughput, base calling accuracy, and assembly algorithms, have increased the utility of Nanopore-sequenced long reads alone, to generate high-quality genome assemblies. Additionally, our study shows that a combined approach to genome annotation employing both transcriptomic- and orthology-based methods can improve the completeness of genome annotations. By providing both a resource for future analyses on an important solitary bee species but also novel, cost-effective approaches to genome assembly and annotation, we aim to contribute to an urgently needed, growing research interest in the genomics of understudied pollinators.

## Materials and Methods

### Sample collection

For the purpose of genome assembly, in March 2023, one male adult was collected and immediately snap-frozen after emergence from pre-placed nest boxes from Rottleberode (Südharz, Saxony-Anhalt, Germany, 51°31′N, 10°57′E). For the purposes of gene model prediction, we sampled from adult and developmental stages to increase gene discovery rates, as certain genes may only be expressed during certain stages. For this, we collected the following: 1) one female adult was sampled from the same nest box as the male used for genome assembly and immediately snap-frozen after emergence; and 2) male and female pupae were sampled in 2023 and 2024 from additional pre-placed nest boxes, and immediately snap frozen. All samples were stored at −80°C until DNA and RNA extractions were performed, respectively. Lastly, for the purposes of examining intraspecific genetic variation present within a representative *O. cornuta* population, we sampled female adults from 10 independent sites (more than 50km apart) across Germany (for exact locations see Supplementary Table 1) for whole-genome resequencing (n=10 individuals total) and downstream population genetic-based analyses. For sampled individuals, field-based species identification was initially performed based on morphological features (Westrich 2019), before bees were immediately conserved in absolute ethanol in the field with subsequent storage at −80°C. Species identification for all samples was later assessed and validated using computational techniques (Supplementary Information: Computational validation of species identification).

### DNA extraction and long-read sequencing

High-molecular-weight DNA was extracted from the thorax of the single adult male using the New England Biosystems Monarch HMW DNA extraction kit, yielding 7.5µg of DNA (260/280nm ratio of 1.7, 230/280nm ratio of 1.0, at a concentration of 150ng/µl according to a Nanodrop ND-1000). A Nanopore sequencing library was generated using a ligation-based kit (SQK-LSK114) from Oxford Nanopore Technologies (ONT). Sequencing was performed on approximately half of the capacity of a PromethION flow cell (v10.4.1 FLO-PRO114M) in two runs of 20 hours and 48 hours, respectively, with each run separated by a flow cell wash. The initial 39.57Gb of raw data (4.86 million reads at N50 of 17kb) were subjected to basecalling a second time with dorado (v0.7.0+71cc744, model sup5.0.0) on a Geforce GTX3090 GPU. Reads were then filtered by quality (≥Q90) and length (≥8kb) (Filtlong v0.2.1, seqtk v1.4-r122), and, additionally, error-corrected with HERRO model_v0.1 for R10.4.1 data (Stanojevic et al. 2024) after which reads less than 5kb were removed.

### RNA extractions for transcriptome-facilitated genome annotation

For the prediction of gene models during genome annotation, we generated a transcriptomic dataset for the head tissues of a single *O. cornuta* adult female. To achieve this, from a snap frozen sample, we removed the head from the thorax, and transferred it to a 1.5ml centrifuge tube containing 200μl of TRIzol. The sample was manually homogenised using a micropestle until the head was visually disrupted. Total RNA was subsequently extracted using a TRIzol-chloroform-based extraction protocol (T. J. Colgan et al. 2011), and then purified using a Monarch RNA purification kit (NEB) before being DNase-treated (TURBO DNase, Qiagen) to remove putative DNA contaminants. RNA was quantified and assessed using the FastGene NanoSpec Photometer (FG-NP01), before being stored at −80°C. DNase-treated RNA was shipped on dry ice to a commercial supplier (Novogene, Germany) where quality was further assessed using an Agilent Bioanalyzer. An mRNA-enriched library was prepared and paired-end sequencing (2*150bp) was performed to produce a minimum of 50 million paired-end reads. For the additional pupae samples, total RNA was extracted from four different leg tissues (hind tibia, mid tibia, hind basitarsus, mid basitarsus) from 30 samples (15 female, 15 male) using mechanical homogenization (Tissue Lyser LT, Qiagen) and a Zymo DirectZol kit, which included DNAse digestion. For the female adult, a mRNA-enriched library was prepared, while for the pupae samples mRNA was extracted from total RNA (poly-T oligo-attached magnetic beads), followed by fragmentation and cDNA synthesis with random hexamer primers. Sequencing (paired end 150 bp) was performed by Novogene (Germany) generating a mean read count of 55,206,130 across samples (min: 12,042,090, max: 162,207,301).

### DNA extractions for population genetic analyses

For the population genetic-based analyses, we removed ethanol-preserved samples (n=10) from the −80°C freezer. As abdomens can harbour microbial contaminants, we removed them using microscissors and the remaining tissue was subsequently washed with three rounds of autoclaved phosphate-buffered saline (PBS) to remove residual ethanol. Tissue homogenisation was performed using micropestles on tissue samples in 1.5ml Eppendorf tubes filled with 200µl of PBS. Following this, DNA was extracted using the Monarch Genomic DNA Purification kit, according to the manufacturer’s instructions. After extraction, the quality and quantity of isolated genomic DNA were checked using NanoDrop and Qubit photometric measurements, as well as by using low concentration agarose gel electrophoresis at low voltage. Only samples with a minimum total DNA amount of 1.2µg, 260nm:280nm absorbance spectrum ratios of at least 1.8, and visible sharp bands in electrophoresis gels were considered for sequencing. Extracted DNA was sent to a commercial sequencing company (Novogene, UK) where further quality assessments using an Agilent Bioanalyzer were performed. After quality checks, for each sample, an individual PCR-free library was produced using a NEB library kit, individually barcoded, and sequenced on an Illumina NovaSeq 6000 (2*150bp) to a minimum sequencing output of 3Gb per sample, resulting in a mean read count of 32,695,578 across samples (min: 26,511,816, max: 38,517,412, Supplementary Table 1), as well as a mean sequencing depth of 19x, based on the total length of chromosomal scaffolds.

### Genome assembly

Using a combination of filtered long FASTQ reads and HERRO-corrected FASTA reads, we generated and assessed different *de novo* assemblies using flye v2.9.3 (Kolmogorov et al. 2019), specifically, generating assemblies with variation in minimum overlap (10,000, 8,000, and 6,000bp) with presets for pacbio-hifi (corrected reads), nano-corr (corrected reads), and nano-hq (un-corrected reads). As a complementary approach, we generated a second set of assemblies using hifiasm v0.24.0 (Cheng et al. 2024) using the “–ont” flag with uncorrected reads and with five different parameter sets (see code on GitHub). From each resulting assembly, we removed contigs shorter than 5kb using seqtk as long reads used for the assemblies were either 8kb or longer meaning that contigs shorter than this length represent potential sequencing artefacts or are biologically less informative. Each assembly was assessed for summary statistics (assembly_stats, https://github.com/sanger-pathogens/assembly-stats), gene completeness (Compleasm v0.2.7 with database hymenoptera_odb12) and the presence of the canonical hymenopteran telomere repeat “AACCT” using tidk, (Brown, Manuel Gonzalez de La Rosa, and Blaxter 2025).

The best assembly was assessed and chosen according to the lowest number of missing or duplicated BUSCO genes, the highest number of identified telomeric regions, and the longest contigs. Based on these criteria, “hifiasm3” was selected as the best assembly, which was generated using hifiasm with the following parameters: “--telo-m AACCT --primary -l 2 -s 0.65 -D 10.0 -N 300 --dual-scaf --ont”. This assembly was polished using medaka v1.12.1 (https://github.com/nanoporetech/medaka), followed by the purging of haplotigs and overlaps using purge_dups v.1.2.5 (Guan et al. 2020), as well as the sorting by length in a descending order (i.e., longest to shortest) and renaming of contigs using seqkit v2.5.1 (Shen et al. 2016). The quality of the assembly was then again assessed using assembly_stats, Compleasm, and tidk, which collectively provide information on basic summary statistics of genome size and structure, assembly completeness, and potential contaminants. As an additional step, assembly completeness relative to the long reads used for assembly was assessed with merqury (2022-09-07, (Rhie et al. 2020)).

The putative mitochondrion was assembled in a separate process. First, we used BLASTN with a set of bee mitogenomes to identify six contigs likely of mitochondrial origin. One of these contigs, representing a putative single mitochondrial genome copy, was selected as a reference so as to allow extraction of Nanopore long reads that matched it after mapping with minimap2 (v2.28-r1209). These extracted reads were subsequently assembled (flye v2.9.3-b1797) into a single contig of 35,525bp. This assigned mitogenome was then annotated with mitos v2 (Donath et al. 2019).

### Assessment of possible contamination

We screened the assembled nuclear genome using BlobToolKit (Challis et al. 2020) to identify scaffolds of unusual GC content-to-read coverage ratios, as well as scaffolds assigned to non-target organisms, which likely represent contaminants. In addition, the assembly was further assessed for contamination using the NCBI Foreign Contamination Screen (fcs v0.4.0) pipeline with NCBI’s *Osmia cornuta* TaxID 185587 specifically selected.

### Repeat annotation

Putative repetitive elements in the genome assembly were identified and annotated using EarlGrey v5.1.1, which incorporates RepeatMasker 4.15 with the DFAM 3.7 and RepBase (2018) databases, and was run with clustering and softmasked output. This soft-masked genome assembly was used as the input for gene model predictions (**‘**Gene annotation’). Satellites were identified with Satellite Repeat Finder (https://github.com/lh3/srf), while microsatellites and regions of low complexity were identified using a combination of the following complementary tools: etrf (-m 200 -l 13; https://github.com/lh3/etrf), trf-mod v4.10.0, SA-SSR v0.1 alpha (--min-repeats 5 --min-nucs 10 --max-ssr-len 12 --max-seq-len 25000000), ULTRA v1.0.0 beta 12 (--min_unit 4 --min_length 10 --period 300, filtered for minimum length of 100bp), tantan v49 (-f4, filtered for minimum repeat number=5), sdust (-w 64 -t 20, filtered for minimum length=10), and hrun (filtered for minimum length=10, seqtk v1.3-r115-dirty). Low mappability regions were identified using genmap v1.3.0 (parameters: -K 35 -E 2), with subsequent filtering based on mappability below 0.05 and a minimum length of 100bp. From the EarlGrey annotations, we removed entries classified as “Unknown” and “Satellite” to create a set of transposable element (TE) annotations. Putative microsatellite and low complexity regions were merged and overlaps with TE annotations removed using bedtools to create a set of repeat annotations.

### Gene annotation

To increase our ability at predicting gene models for tissue-specific or lowly-expressed genes, we complemented our samples of adult head (n=1) and pupal leg (n=120) tissue with publicly available samples of antennal tissue (n=8) generated by the Institute of Applied Genomics (NCBI BioProject: PRJNA185987), as well as a publicly available whole-bodied sample generated through the 1KITE project (n=1, NCBI BioProject: PRJNA252331). Prior to gene model prediction, we performed data quality evaluations for all samples. For each sample, we first examined quality metrics generated using FastQC (Andrews and Others 2010), which provides information on base quality, as well as presence and extent of adapter contamination. As a complementary approach, we investigated the proportion of raw reads mapping to the generated genome assembly using HISAT v2.2.1 (Kim et al. 2019), finding high mapping rates (mean: 96.06%). We then combined and visualised the outputs for both sets of quality assessments using MultiQC v.1.29 (Ewels et al. 2016). Based on the results of the quality assessment, we removed adapter sequences, which can affect mapping rates and quality, and filtered reads by quality (PHRED score ≥ 15) and length (minimum length ≥ 50 bp) using fastp v.0.20.1 (Chen et al. 2018).

The filtered reads were then used in conjunction with a protein database comprising arthropod-derived proteins from OrthoDB v12 (Kuznetsov et al. 2023), published hymenopteran proteins (Kapheim et al. 2015), and all proteins from the curated Swiss-Prot database (UniProt Consortium 2023) to predict gene models within the soft-masked genome assembly using BRAKER3 v3.0.8 (Gabriel et al. 2024). Initial predictions from this approach were further processed to identify and correct putative incorrect gene models, such as by splitting up gene models with no overlap among their transcript “children” into separate gene models, and by removing transcripts with internal stop codons or missing start or stop codons. As a complementary approach, gene models were independently predicted via a second approach based upon orthology to the *O. bicornis* reference genome (iOsmBic2.1, GCF_907164935.1) using TOGA v1.1.7 (Kirilenko et al. 2023). The resulting predictions, based upon orthologous matches, were filtered to include only predictions classified as “intact”, “partially intact”, or “uncertain loss”. Genes with no overlap among any of their transcripts were split into separate genes. During gene model prediction, when transcripts from *O. bicornis* were aligned to the *O. cornuta* genome, some transcripts mapped to different chromosomal scaffolds or contigs than the rest of the transcripts from the same source gene. These inconsistently mapped transcripts were removed to make sure that each newly predicted gene model in *O. cornuta* was single-locus. As the orthology approach to gene prediction sometimes generated very short introns of 1-3bp in otherwise correctly predicted transcripts, we joined adjacent exons if they were separated by less than 4bp. Incomplete gene predictions with missing start or stop codons, or internal stop codons, were removed to retain only the highest quality predictions. We then computed overlap statistics between the two sets of annotations and complemented the BRAKER3 gene models with any additional transcript isoforms that were unique to the filtered TOGA predictions.

We inferred functional annotation from our translated, predicted proteins via homology and structural similarity to the publicly available annotation databases InterPro, Egg-Nog, BUSCO, dbCAN, SignalP, and UniProt (Blum et al. 2021; Huerta-Cepas et al. 2019; Simão et al. 2015; Zheng et al. 2023; Almagro Armenteros et al. 2019; UniProt Consortium 2025) using Funannotate v1.8.17 (Palmer and Stajich 2020). Lastly, we quality checked our final genome annotation using Compleasm in “protein” mode (Nevers et al. 2025; Manni et al. 2021) and OMark v0.3.1 (Nevers et al. 2025). To complement our annotation with tRNA and rRNA gene models, we ran tRNAscan-SE (v2.0.12, (Chan et al. 2021) and Barrnap (v0.8, eukaryotic mode: -k euk, (Seemann, n.d.), respectively. Finally, putative mitochondrial fragments integrated into the genome (i.e., NUMTs) were assessed by querying the *O. cornuta* nuclear genome assembly using BLASTN with the putative mitochondrial genome sequence previously identified in our assembly.

### Methylation analysis

As methylation can play an important role in gene regulation, we examined our Nanopore data for base modifications to identify presence and extent of methylation patterns in the O. cornuta genome. We used dorado (v0.9.0) to basecall the raw Nanopore data (pod5) with the base modifications 5mCG, 5hmCG, 6mA, 5mC, and 5hmC, using the super accurate model (dna_r10.4.1_e8.2_400bps_sup@v4.3.0) on a Geforce RTX3090 (with parameter --modified-bases-threshold 0.30). We extracted base modification signals using modkit (v0.4.1) (pileup) from the resulting sorted BAM file. To assess how methylation patterns differed among genomic regions and features (e.g., exonic, intronic, intergenic), we first counted, for each of these feature types, the total number of sites that could potentially be methylated using bedtools intersect. Similarly, we then counted for each feature type, the number of sites that were either weakly methylated (sites covered by at least one read containing a methylated base) or highly methylated (sites with at least 20% of reads containing a methylated base). To account for large differences in the mean length of features, we summarised all counts in 100kb windows (bedtools map -o sum) across the genome, before dividing counts of weakly and highly methylated sites by the total number of sites that could potentially be methylated to generate percentages of methylation across all feature types. Throughout this approach, we processed 5mCG and 5hmCG methylation signals separately.

### Analysis of synteny with *O. bicornis*

To further assess contiguity and completeness of our genome assembly, we examined collinearity, particularly synteny of chromosomal segments, between our *O. cornuta* genome assembly and a reference genome assembly of the closely related red mason bee, *O. bicornis*. To this end, we extracted the longest predicted protein isoforms from both assemblies and ran a protein-to-protein BLAST (BLASTP) search (Camacho et al. 2009) to identify putative homologues. The result of this BLAST search was then provided as input into MCScanX v1.0.0 (Yupeng Wang et al. 2012), which identified collinear blocks between the two assemblies that were subsequently visualised using the web application SynVisio (Bandi and Gutwin 2020).

### Calling, filtering, and annotation of variants

Using the same quality control (QC) pipeline as that for the RNA-seq data, we assessed the raw sequencing data for the short-read whole-genome resequencing (WGS) samples for read-based quality metrics, as well as read mapping rates against our annotated genome assembly using bwa-mem v2.2.1 (Li 2013). Summarisation of mapping statistics was performed using samtools stats (Danecek et al. 2021) and the generation of overview visualisations using MultiQC. Raw reads were quality filtered and adapters trimmed using fastp with quality (PHRED score ≥ 15) and length (minimum length ≥ 50 bp) filters. For variant calling, filtered reads were first mapped against the genome assembly using bowtie2 v2.5.4 (Langmead and Salzberg 2012), followed by variant calling using the BCFtools v1.21 mpileup-call pipeline (Danecek et al. 2021). Variants were filtered using VCFtools v0.1.17 (Danecek et al. 2011) with the following parameters: removal of indels; minimum and maximum mean read depth filters of 5 and 500, respectively; and a minimum quality score of 40. Additionally, a maximum of 20% of samples (i.e., two individuals) per site was allowed to have missing data. After filtering, the mean and median quality (QUAL) scores were 576.2 and 389, respectively. Filtered variants were used as input to a principal component analysis to investigate the genetic relationship between samples (Supplementary Figure 1). Lastly, variants were annotated with SnpEff v5.2f (Cingolani et al. 2012) using protein-coding sequence predictions from our annotation pipeline.

### Inference of nucleotide diversity and linkage disequilibrium

Filtered variants were indexed using tabix (Li 2011) before running pixy v1.2.11 (Korunes and Samuk 2021) to infer nucleotide diversity π and Watterson’s θ across all chromosomes in both 100,000 and 10,000bp sliding windows. In addition to a sliding window approach, π was inferred collectively for each investigated genomic feature (exon, intron, intergenic regions, repeats and low complexity regions) to understand how different functional regions of the genome differ in nucleotide diversity. We inferred linkage disequilibrium decay curves from filtered variant calls using PopLDDecay v3.43 (Zhang et al. 2019). Inbreeding coefficients for population genomics samples were calculated using VCFtools –het.

### Demography inference

To infer past changes of effective population size, we applied PSMC (Li and Durbin 2011) to sequencing data from the 10 sequenced female adults (Supplementary Table 1). Variants were called and consensus sequences generated employing a pipeline that consisted of samtools v1.08 mpileup, bcftools v1.10.2 call, and vcfutils.pl (the ‘vcf2fq’ pipeline). To improve variant detection in low coverage samples we copied and merged forward and reverse reads. For PSMC, we used parameters informed by related species. Based on estimations from the buff-tailed bumblebee (Apidae: *Bombus terrestris*), a mutation rate (μ) of 3.6 × 10^−9^ (Liu et al. 2017) per base pair per generation was chosen, assuming one generation per year as the species is univoltine (Splitt, Schulz, and Skórka 2022; Tasei and Picart 1973). Additionally, the recombination rate estimates for a closely related solitary bee (Megachilidae: *Megachile rotundata*, (Jones et al. 2019)), which is 1.02 cM/Mb and equivalent to a per base recombination rate of 1.02 × 10^-8^, was used, producing a recombination to mutation rate ratio of 2.5 (-r). We applied a maximum TMRCA of −7 (-t) and the atomic time interval pattern was set to “2+2+25*2+4+6” (-p), to increase comparability (Hilgers et al. 2025). To assess variance in effective population size estimates, we performed 100 bootstrap replicates per individual.

## Data Availability Statement

Data and scripts required for reanalysis are provided as supplementary files and will be made publicly available on GitHub and Zenodo upon acceptance. The genome assembly will be made publicly available on NCBI under the BioProject accession PRJNA1289724 upon publication. DNA resequencing and RNA data will be made available under the BioProject accession PRJNA1289047. Additional RNA data from pupal leg tissue will be made publicly available as hintfile (.gff) via Github and Zenodo upon publication.

## Acknowledgments

We would like to thank the German Federal Environmental Foundation (Deutsche Bundesstiftung Umwelt, DBU) for supporting JSM with a doctoral scholarship (20021/743), and the Eva Crane Trust (ECTA_20210915, United Kingdom), for financial contribution to the *Osmia cornuta* population genomics. As part of the GEvol SPP 2349, this study was funded by the Deutsche Forschungsgemeinschaft (DFG, German Research Foundation) – project number 503360601 (ES, XH, TJC, KN), and the GenEvo Research Training Group (GRK2526) – project number 407023052 (EAB, TJC). We thank Dr. Francieli das Chagas for confirmation on the identity of yellow cluster genes. We would also like to thank the environmental offices of Baden-Württemberg, Bavaria, Berlin, Bremen, Hesse, Mecklenburg-Western Pomerania, Lower Saxony and Saxony, and especially Dr. Peer Schnitter (Landesamt für Umweltschutz (LAU), Saxony-Anhalt), for collection permits. For caretaking of a nesting site, logistics and provision of flower resources, we thank Joachim and Petra Stolle. We also thank the molecular lab at the LIB, in particular Anja Bodenheim, for technical support with ONT sequencing. We would like to thank the HPC facility at JGU for access to computational resources. We also thank the HPC facilities at the LIB for computation support and the LIB for startup-funds for a GPU server (ES). For granting copyright to images of *Osmia cornuta*, we thank Michael and Mandy Fritzsche from Apidarium (www.apidarium.de).

## Author Contributions

ES, JSM, and TJC conceived the study. ES generated the long-read data. JSM and TJC generated whole-genome and RNA-seq data, KN and AH generated RNA-seq data for which ES and XH performed sampling. ES and XH performed the repeat inference. ES performed the methylation detection. ES and JSM did the contamination screening. ES generated the genome assembly. JSM generated the genome annotation. JSM performed synteny and overlap inferences. JSM performed statistical analyses and generated all figures. JSM performed population genetic analyses with EAB contributing the PSMC analysis. JSM wrote the draft, while ES and TJC contributed to the final manuscript. All authors gave final approval for publication.

