## Supplementary Information for "A chromosome-level genome assembly and resequencing data reveal low DNA methylation and reduced diversity in the solitary bee pollinator *Osmia cornuta*"

### Computational validation of species identification

Species identification for all samples was initially performed based on morphological characteristics but later confirmed computationally. To that end, we first identified the locus of the *O. cornuta* cytochrome c oxidase subunit 1 (COX1) sequence within our genome assembly via orthology with the COX1 sequence of the *Osmia bicornis* reference genome (iOsmBic2.1, GCF_907164935.1) using BLASTN (v2.14.1+, (Camacho et al. 2009)). The *O. cornuta* COX1 orthologue sequence was then extracted from this locus using samtools faidx (Danecek et al. 2021) and searched against COX1 sequences housed in the database of Barcode of Life Data Systems (BOLD, queried on the 3rd of October, 2024, (Ratnasingham and Hebert 2007)), validating that the male, which we used to generate the genome assembly was correctly identified as *O. cornuta*. For all other DNA- and RNA-seq samples, reads were mapped against the *O. cornuta* genome assembly using STAR (v2.7.11a, (Dobin et al. 2013)) and for each sample a consensus sequence of matches against the identified *O. cornuta* COX1 locus was constructed using samtools consensus (Danecek et al. 2021). The resulting COX1 sequences were then searched against BOLD using their web API to confirm species identifications.

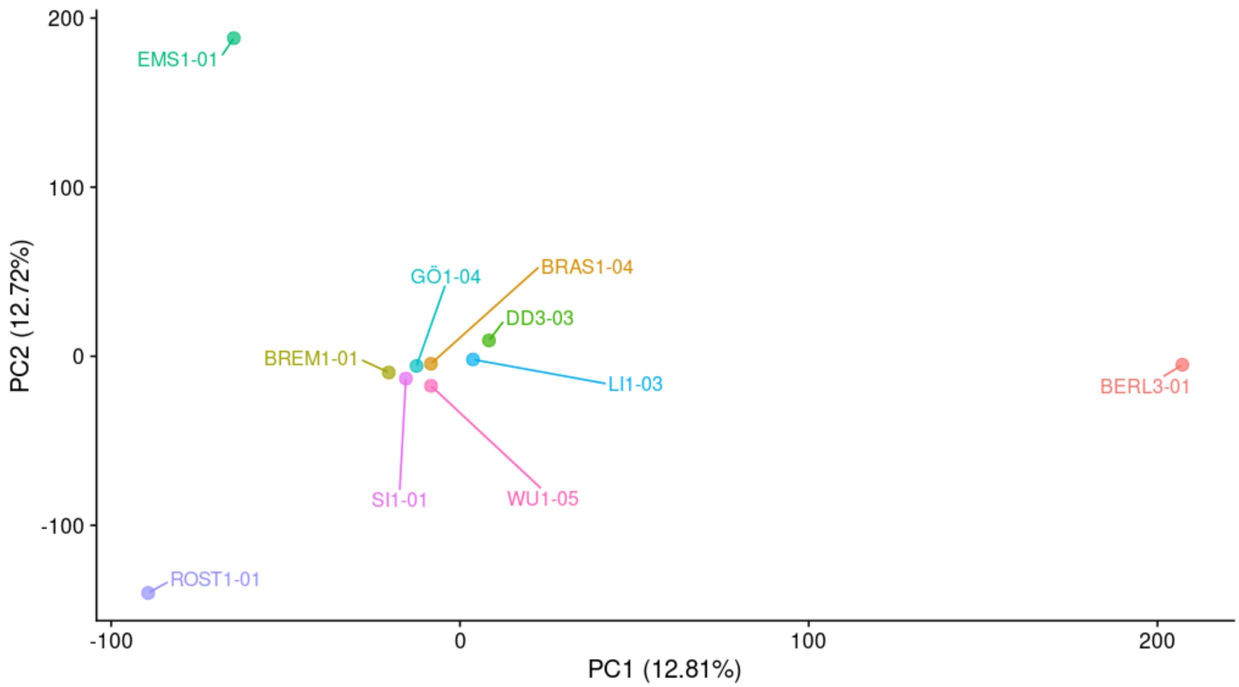


**Supplementary Figure 1. Genetic relationship among sampled *Osmia cornuta*.** Scatterplot displaying the first two principal components calculated from a principal component analysis based on single nucleotide polymorphisms called within a population of *Osmia cornuta* sampled across different locations in Germany. The x-axis shows the first principal component and the total variance explained by it, while the y-axis shows the second principal component and the variance it explains. Each coloured and labeled dot represents a single female *O. cornuta*.

| Sample ID | Sampling Date | Latitude | Longitude | Altitude [m] | Inbreeding Coefficient F | Raw Read Count |
|---|---|---|---|---|---|---|
| BERL3-01 | 12/04/23 | 52.60 | 13.40 | 49 | 0.21189 | 38,517,412 |
| BRAS1-04 | 29/03/23 | 52.31 | 10.52 | 69 | 0.10376 | 28,428,320 |
| BREM1-01 | 10/04/23 | 53.07 | 8.80 | 8 | 0.19567 | 29,153,738 |
| DD3-03 | 05/05/22 | 50.88 | 13.68 | 387 | 0.18829 | 36,106,378 |
| EMS1-01 | 10/04/23 | 52.36 | 7.30 | 41 | 0.23890 | 36,025,098 |
| GÖ1-04 | 23/04/22 | 48.71 | 9.63 | 338 | 0.11099 | 26,511,816 |
| LI1-03 | 02/05/22 | 50.39 | 8.05 | 129 | 0.12830 | 31,449,700 |
| ROST1-01 | 11/04/23 | 54.09 | 12.09 | 26 | 0.19660 | 34,304,438 |
| SI1-01 | 23/04/22 | 49.22 | 8.79 | 197 | 0.04068 | 31,255,604 |
| WU1-05 | 27/04/22 | 49.77 | 9.93 | 233 | 0.11870 | 35,203,272 |

**Supplementary Table 1. Sample site information for *Osmia cornuta* used for population genomics.** The table is showing for each *O. cornuta* that was considered for the population genomics analyses, the sample identifier, the date of sampling, sampling site coordinates, inbreeding estimates calculated from expected versus observed homozygosity, and the total count of raw reads generated for each sample.
